# A covalently crosslinked bioink for multi-materials drop-on-demand 3D bioprinting of three-dimensional cell cultures

**DOI:** 10.1101/2021.02.18.431759

**Authors:** Robert H. Utama, Vincent T. G. Tan, Kristel C. Tjandra, Andrew Sexton, Duyen H. T. Nguyen, Aidan P. O’Mahony, Julio C. C. Ribeiro, Maria Kavallaris, J. Justin Gooding

## Abstract

In vitro three-dimensional (3D) cell models have been accepted to better recapitulate aspects of in vivo organ environment than 2D cell culture. Currently, the production of these complex in vitro 3D cell models with multiple cell types and microenvironments remains challenging and prone to human error. Here we report a versatile bioink comprised of a 4-arm PEG based polymer with distal maleimide derivatives as the main ink component and a bis-thiol species as the activator that crosslinks the polymer to form the hydrogel in less than a second. The rapid gelation makes the polymer system compatible with 3D bioprinting. The ink is combined with a drop-on-demand 3D bioprinting platform consisting of eight independently addressable nozzles and high-throughput printing logic for creating complex 3D cell culture models. The combination of multiple nozzles and fast printing logic enables the rapid preparation of many complex 3D structures comprising multiple hydrogel environments in the one structure in a standard 96-well plate format. The platform compatibility for biological applications was validated using pancreatic ductal adenocarcinoma cancer (PDAC) cells with their phenotypic responses controlled by tuning the hydrogel microenvironment.

## 1. Introduction

*In vitro* cell culture has been used in routine cancer biology research as a tool to study cellular mechanisms. Conventionally, *in vitro* cell culture is predominantly performed on a flat, two-dimensional, monolayer format. More recently, the *in vitro* cell culture paradigm has begun to shift towards three-dimensional (3D) culture formats where the cell-to-cell and cell-to-matrix interaction can be better recapitulated.^[1,2]^ However, widespread uptake of 3D cell models is encumbered by the challenge of producing 3D cell cultures with sufficient throughput and reproducibility.^[3]^ Furthermore, the creation of complex 3D cell models comprising multiple cell types in different extracellular matrix (ECM) environment ^[4,5]^ are exceedingly difficult to achieve due to the lack of availability of a modular ECM mimic system and an accurate method to position different mimics and/or different cell types at defined locations to create such models.

3D bioprinting has the potential to creates structurally complex scaffolds.^[6,7]^ Extrusion-based bioprinting has recently found extensive applications in tissue engineering, where it benefits from the on-demand prototyping capability for few large, multi-material constructs.^[8-10]^ In contrast to tissue engineering, bioprinter for generating complex 3D *in vitro* cell models must exhibit the capability to print multiple materials and cell types into a 3D structure in high throughput.^[11]^ Drop-on-demand bioprinting is a suitable candidate for this application. Examples exist in the literature of drop-on-demand bioprinters show the capability of printing multiple materials in the one structure.^[12,13]^ We recently reported a bespoke drop-on-demand bioprinting platform for rapid production of 3D *in vitro* cell models.^[14]^ Drop-on-demand bioprinters however are limited by the requirement for low cell concentration, low viscosity bioinks and rapidly-gelating hydrogel systems.^[15]^

Of the different hydrogel systems available, synthetic hydrogels as ECM mimics possess much lower batch-to-batch variation in properties compared with biologically derived ECMs. Furthermore, synthetic hydrogels offer unprecedented control over the physical and adhesive cues presented to the cells.^[16,17]^ In particular, poly(ethylene glycol) (PEG)-based hydrogels stand out as a highly modifiable material that can be adjusted to induce specific biological responses.^[18,19]^ To overcome the biologically inert nature of synthetic hydrogels, peptide sequences are commonly incorporated to the hydrogel to induce cell adhesion, using motifs such as RGD^[20]^ and IKVAV,^[21]^ and cell motility (using matrix metalloproteinase (MMP)-responsive peptide sequences).^[22]^ For a polymer system to be compatible with scaffold-free, drop-on-demand 3D bioprinting, it requires the bioink to gel rapidly within a few second after printing.^[15]^

The purpose of this paper is to explore a new bioink for a modular drop-on-demand bioprinting platform for multi-material printing of complex 3D structures in high-throughput that covalently crosslinks. The ink is based on a combination of PEG-hydrogels with Michael addition reactions between thiols and maleimide moieties as this system offers a hydrogel that has tunable properties, low viscosity gel precursors and potentially rapid gelation at neutral pH.^[23,24]^ The main ink component is a 4-arm PEG based polymer with distal maleimide derivatives at the end of each arm (PEG-4MAL) and a bis-thiol activator that cross-links the polymer to form the hydrogel. We show that this reaction is sufficiently rapid to sustain drop-on-demand 3D bioprinting via a covalent linkage between the polymer that forms the bulk of the ink and the crosslinking activator. The modularity and compatibility of the PEG hydrogel scaffold for cell research was demonstrated against pancreatic ductal adenocarcinoma (PDAC) cancer cells which showed different phenotypic responses in different ECM mimic environments.^[25]^ This is a significant advance on our previous report on the bespoke bioprinter that is capable of bioprinting viable cells from a 250 million cells/mL cell suspension and producing a full 96-well plate of single spheroid embedded in an alginate scaffold in under 80 minutes.^[14]^ A combination of the PEG-hydrogel, thiol-Michael addition reaction and the bespoke bioprinter enabled us to create a 3D structure comprising multiple different hydrogel environments that are compatible with *in vitro* 3D cell as**s**ays.

## 2. Hydrogel Formulation and 3D Bioprinting Optimization

We previously reported a 3D bioprinting platform designed for the high-throughput production of 3D spheroids. To enable the bioprinting of multiple cells or biomaterials, the bioprinter was upgraded with a sterile bioprinting cartridge that carries multiple functional inks (Figure 1a-i). and a printhead with eight independently-addressable nozzles for printing complex models (Figure 1a-ii) for printing in a multi-well plate (Figure 1a-iii). The bioprinter utilizes a fly-by printing logic that enable high-throughput production of *in vitro* 3D cell models. During fly-by printing, the ejection of ink droplets occurs as the printhead is travelling on the x-axis at a constant velocity. Firstly, the maleimide-containing bioink droplets were printed across the full row of the well plate as the printhead travels on the positive x-axis direction (Figure 1b-i). On the return flight, thiol-containing activator droplets were printed on top of the bioink droplets to initiate the hydrogel formation (Figure 1b-ii). Upon contact, gelation occurred instantaneously via a nucleophilic addition reaction of the thiolates onto the maleimide group (Figure S1b-c, (Supporting Information)). The process of printing a row of hydrogel in a 96-well plate, at a given y position, was completed within 5 seconds.

**Figure 1.**
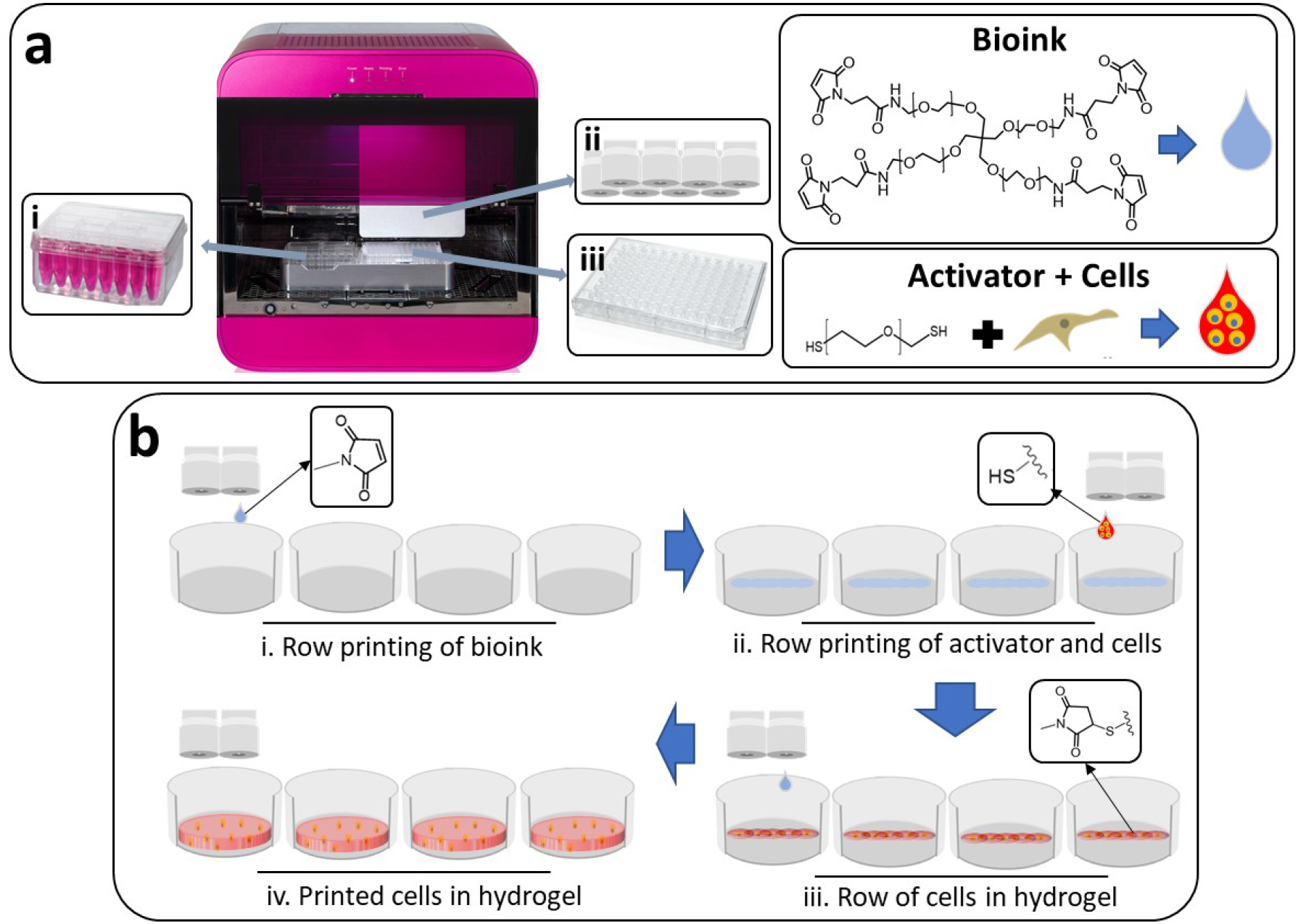
3D bioprinting process based on drop-on-demand printhead. (a) A standalone RASTRUM 3D bioprinting platform (Inventia Life Science) with an integrated biosafety cabinet. (a-i) PEG-4MAL bioinks and cells suspended in bis-thiol activators are prepared in RASTRUM cartridge. (a-ii) Bioinks and activators are then primed automatically into the printhead, equipped with 8 independently addressable nozzles. (a-iii) Printing is done into a standard multi-well plate, such as a 96-well plate. (b) Schematic of the fly-by bioprinting process.

The hydrogel system comprised a 4-arm poly(ethylene glycol) maleimide (PEG-4MAL) bioink, with or without thiolated adhesion peptides, and either bisthiol-PEG or MMP-responsive activator which crosslinks the polymer into the hydrogel (Figure 1a and Figure S1a (Supporting Information)). To obtain the optimum hydrogel composition for bioprinting, we first studied the effect of bioink concentration and the molar ratio reactive groups on gelation speed in bulk. As the fly-by printing process deposits adjacent hydrogel droplets in sub-second, we hypothesized that the droplet gelation must be within the same time interval to avoid droplets wicking together. To achieve this, we chose pH 7.4 gelation condition to increase the thiol-maleimide reaction kinetics.^[26]^

Gelation was assessed by a tube inversion test, whereby gelation was regarded as complete when no flow of material was observed when a vial containing the gel was turned upside down. Assessment of the gelation time of the manually prepared hydrogels (i.e. without using the printer) as a function of the concentration of the PEG-4MAL bioink showed there was a slowing of the gelation time from instantaneously upon mixing (< 1 second) for 10 wt% to requiring greater than 3 seconds for ink at less than 5 wt% concentration (Table S1, (Supporting Information)). Utilizing the 10 wt% PEG-4MAL bioink, we then assessed the effect of thiol:maleimide concentration on gelation time. At 1:2 thiol:maleimide concentration ratio, a similar instantaneous gelation (< 1 second) to the 1:1 ratio hydrogel was observed. In contrast, a soft gel that flowed under gravity was obtained after 1 minute of incubation when the bis-thiol compound was in excess.

To understand the effect of gelation time on the fidelity of the 3D bioprinted structure, we designed a 3-layer, 5×5-droplet 3D structure, with a 0.75 mm droplet-to-droplet spacing, giving a theoretical dimension of 3.75 x 3.75 mm (LxW) (Figure S2a). Structure fidelity was assessed based on the definition of the shape, the presence of excess liquid as well as droplet satellites around the gel. Prior to bioprinting, the viscosity of the 10 wt% PEG-4MAL bioink and the corresponding bisthiol-PEG activator were confirmed to be suitable for drop-on-demand bioprinting at 4 and 1 cP (**Figure 2**a), respectively.

**Figure 2.**
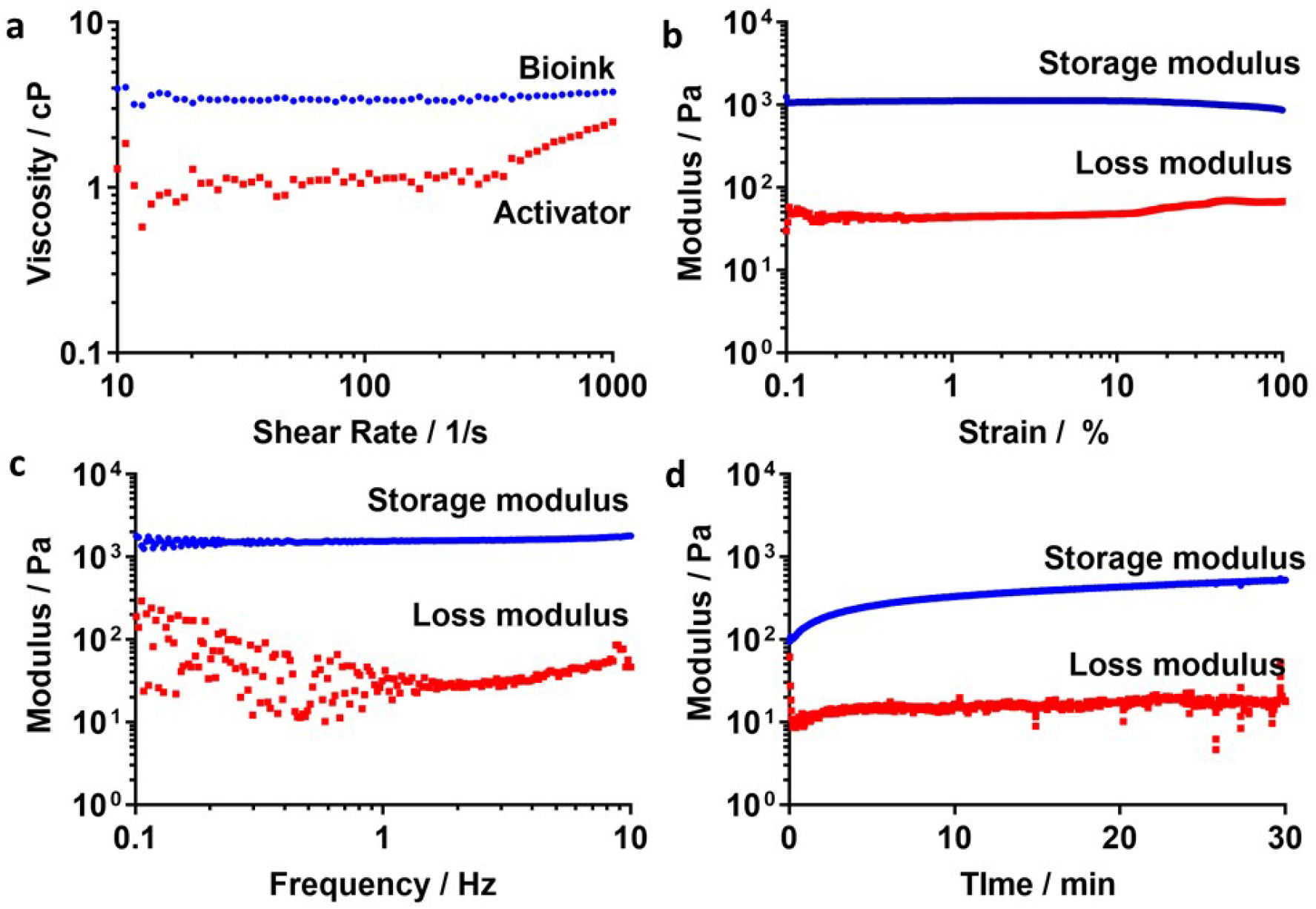
Rheological properties of the developed materials. Mechanical properties of the hydrogel were studied on the PEG-4MAL hydrogel with 1 mM RGD and crosslinked with bisthiol-PEG. (a) Shear-sweep (10 – 1000 1/s) viscosity measurement of the 10 wt% PEG-4MAL bioink and the complementing bis-thiol activator showing a viscosity value of less than 10 cP for both. (b) The frequency and (c) strain sweep studies of the 5 wt% PEG hydrogel to determine the constant strain and frequency values within the LVR region. (d) 5 wt% PEG hydrogel stiffness analysis was conducted at constant frequency and strain values within the viscoelastic region of the hydrogel, showing a stiffness value of around 0.53 kPa.

Bioprinting using 5 wt% PEG-4MAL bioink did not successfully reproduced the designed cuboid structure, with no discernible straight edges, in all wells. Satellite droplets around the structure (red circle) were observed, which were attributed to splashing when the printed droplet hit a non-gelled surface (Figure S2b (Supporting Information)). In contrast, printing with the 10 wt% PEG-4MAL bioink produced a distinct rectangular shape with straight edges, with no droplet satellites, in all wells (Figure S2b (Supporting Information)). The length was measured to be ∼4% larger than the theoretical calculation at 3.92 ± 0.07 mm, while the width was ∼8% smaller at 3.45 ± 0.07 mm. The reduction in structure width is attributed to utilized printing process of building the 3D structure line-by-line, whereby a small degree of wicking to the preceding line occurs when a subsequent line is printed, thus reducing the width of the structure.

We also tested the effect of the thiol-maleimide concentration ratio imbalance on the printed structure fidelity (Table S2, (Supporting Information)). Incorrect structural definition relative to the designed rectangular shape was observed when excess thiol was present (Figure S2c, (Supporting Information)). This was due to incomplete gelation; hypothesized to be a result of many of the bisthiol activators only being conjugated at end single to available maleimide groups rather than both thiols on a given activator molecule being involved in cross-linking between the PEG-MAL chains. Interestingly for the hydrogel with excess maleimide groups, the structure was not as well-defined (indicated by the more rounded vertices) as the hydrogel formed from a 1:1 ratio of maleimides of thiol moieties, even though both samples achieved instantaneous gelation (Figure S2c (Supporting Information)). We attributed this to the insufficient thiols within the hydrogel system to achieve a crosslinking density for the hydrogel to be able to mechanically support the 3D structure without collapsing. Based on the results, we concluded that a bioink at 10 wt% or above PEG-4MAL concentration printed at 1:1 reactive group ratio is needed to bioprint a 3D structure consistently.

Having selected 10 wt% PEG-4MAL 20 kDa bioink as the suitable concentration for bioprinting, we then assessed the rheological properties of the corresponding 5 wt% PEG hydrogel generated by mixing the bioink and the activator at 1:1 v/v ratio. The hydrogel crosslinked with a bis-thiol crosslinker was used as the model hydrogel for this study. Both frequency and strain sweep measurements were conducted to determine the linear viscoelastic region of the hydrogel to measure the equilibrium gel strength.^[27]^ PEG-4MAL hydrogel showed a linear behavior of constant values as the strain was swept (Figure 2b). A 10 % strain was chosen to ensure sufficiently high signal for subsequent tests. Similarly, the frequency sweep also returned a linear behavior, indicating that the materials remain viscoelastic throughout the chosen frequency range (Figure 2c). As such, 10 Hz frequency was chosen to ensure strong signal from the measurements. Time sweep experiment using 10 % strain and 10 Hz frequency showed no crossover between the storage modulus (*G’*) and loss modulus (*G”*), confirming the instantaneous gelation nature of the system. The crosslinking was found to continue for another 20 minutes before reaching a plateau. The stable storage modulus (*G’*) of the 10 wt% bioink hydrogel was around 0.53 kPa (Figure 2d), equivalent to a Young’s modulus (*E*) value of around 1.5 kPa as calculated according to the formula reported previously^[28]^ with a Poisson ratio (*ν*) of 0.45.^[29]^ The Young’s modulus value of this formulation was confirmed to be suitable for intended application with PDAC cells, according to the previously reported modulus of PDAC of around 1.3 kPa.^[30]^

After confirming the printability of the bioink and the activator to form hydrogel, we then proceeded to show the bioprinter capability in printing structures comprising multiple different materials. To show that different hydrogel compositions could be printed into a 3D structure, we utilized bioinks mixed with red, green or blue colored dyes to represent different hydrogel compositions. Various structure definitions utilizing all three colors were designed, including stripes, concentric squares and checker boxes. Into a 96-well plate, all five 3D structures designed were printed simultaneously within one print-job. Each structure was repeated four times to show reproducibility. The bioprinted hydrogels successfully replicate the designed structures, with a resolution of about 0.5 mm between adjacent droplet of gels bearing different compositions (**Figure 3**). Each repeat also replicated the structure definition, location and fidelity accurately, confirming the high well-to-well reproducibility of the bioprinting process (Figure S3 (Supporting Information)).

**Figure 3.**
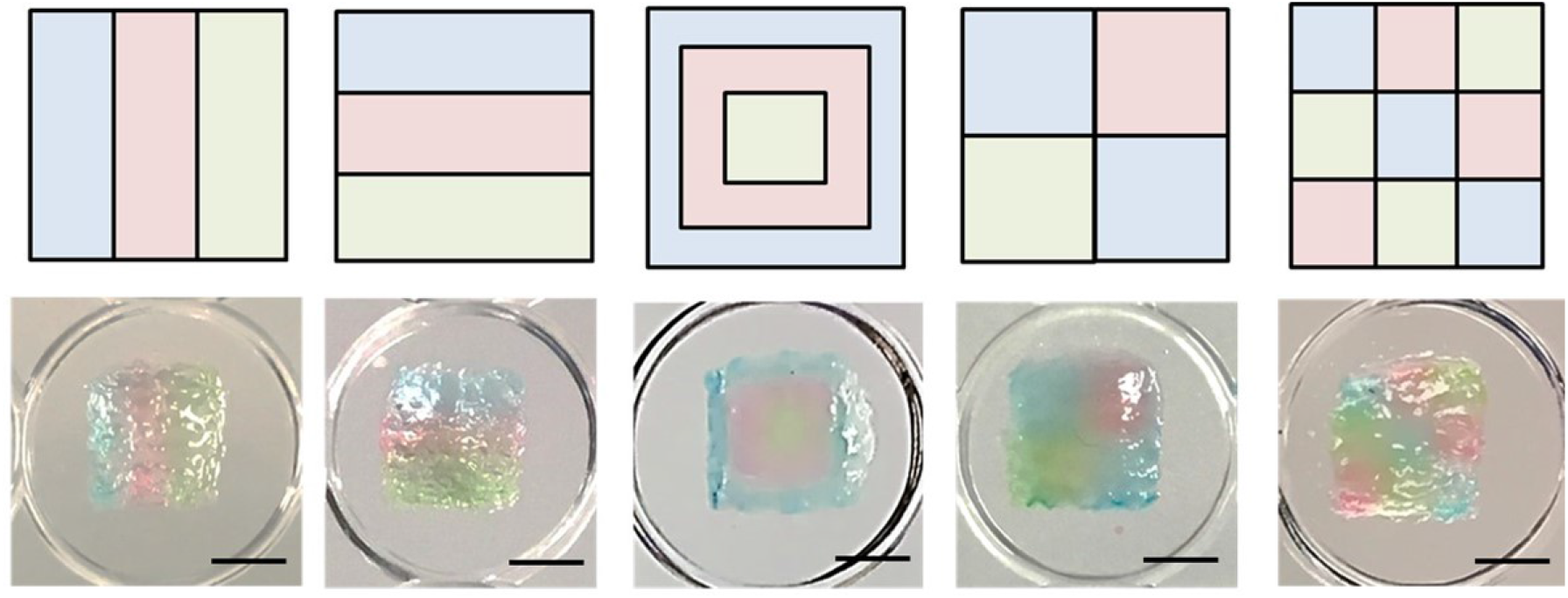
3D bioprinting of structures comprising multiple different materials using colored 4-arms PEG hydrogel and the multiple nozzles available in RASTRUM. Five structures were printed onto each well of a 96-well plate with a diameter of 6.4 mm. The resulting hydrogel structure demonstrates the high resolution and accuracy printing. Scale bars = 2 mm

## 3. Biocompatibility Assessment of the 3D Bioprinting Process

We next investigated the compatibility of the bioinks and the activator with cells. Various factors could affect cell viability during bioprinting process. Each of these were studied independently. First, we studied the effect of the bioink and activator formulation on cell viability. To bioprint 3D encapsulated cells, single cells must initially be mixed with either the bioink or the activator. For the designed PDAC 3D cell models used in this study, we calculated that the cells would need to be suspended in the bioink or activator for ∼60 minutes to print a full 96-well plate. We therefore studied the effect of suspending cells in various PEG-4MAL 20 kDa bioink and activator concentrations (Table S3 (Supporting Information)) for 60 minutes. Cell viability studied using an alamarBlue assay, showed > 92% cell viability when suspended in PEG-4MAL 20 kDa bioink for a hydrogel concentration up to 10 wt% concentration and decreasing to < 50% when cells were dispersed in the bioink for hydrogels at 15 wt% or above (**Figure 4**a). In contrast, cells maintained > 95% viability in the activators for up to 15 wt% hydrogel before dropping to ∼ 90% in the activator for 20 wt% hydrogel. (Figure 4a). Therefore, we tend to suspend the cells in the activator for the bioprinting process. We also tested the effect of printing pressure on cell viability, as shear stress has been previously reported to affect cell viability. Due to the low viscosity nature of the activator, we were able to print 10 million cells/mL suspended in the activator for 5 wt% hydrogel through a ∼200 µm orifice at low printing pressure. Similar cell viability of greater than 95% to the non-printed cells was maintained on cells printed at up to 100 kPa, confirming the gentleness of the printing process towards the cells (Figure 4b).

**Figure 4.**
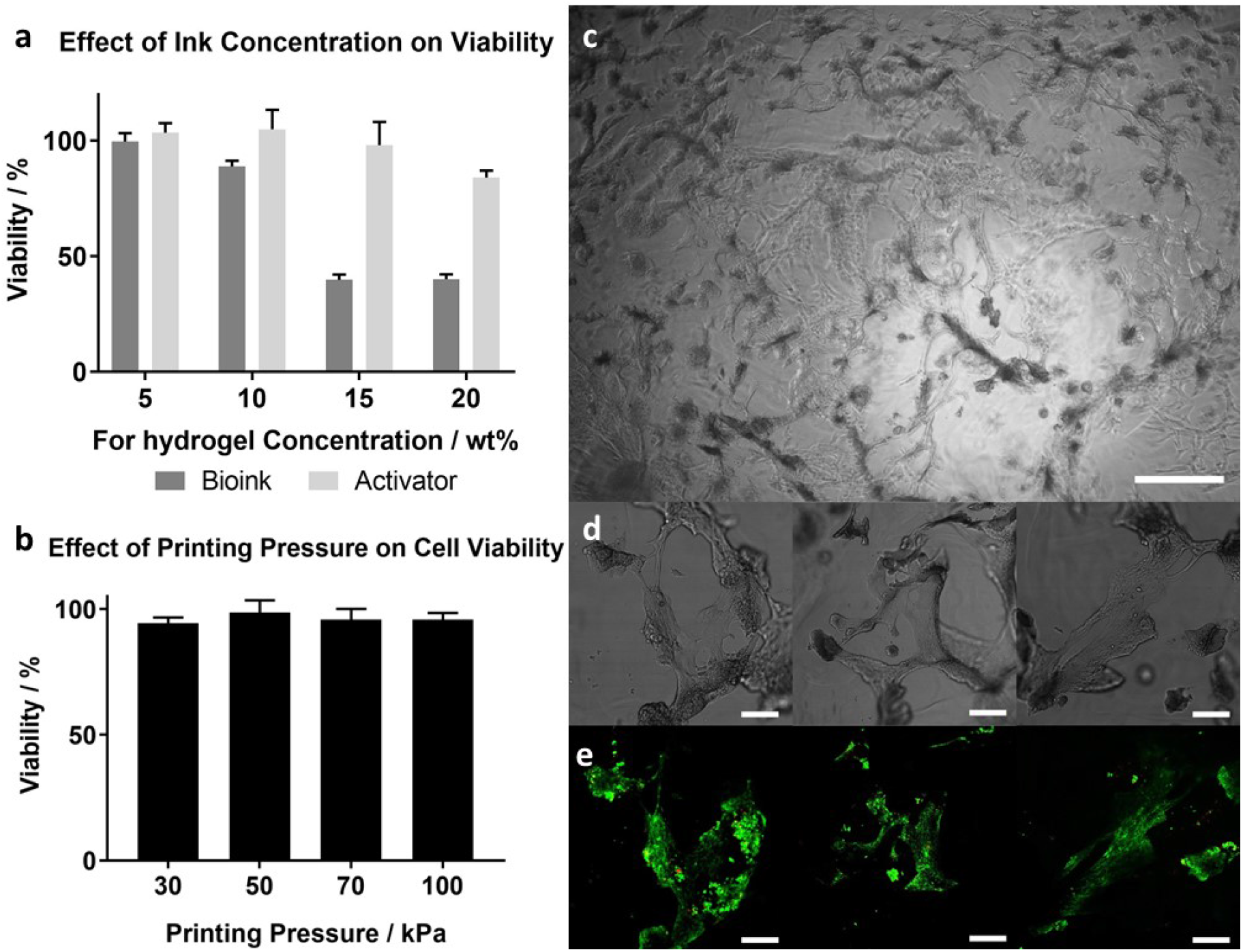
Cell viability studies of pancreatic ductal adenocarcinoma cells when exposed to various 3D bioprinting parameters. (a) AlamarBlue assay on cells suspended in various concentrations of bioink and activator at 10 million cells/mL for 60 minutes, imitating the conditions that the cells are exposed to during the bioprinting process. The study showed the activator has negligible effect on cell viability. (b) AlamarBlue viability assay on cells printed at different printing pressure, showing no visible effect of shear rate on cell viability when compared to non-printed cells. (c) A representative bright field image of pancreatic cells seeded on top of a 3D bioprinted MMP-responsive, PEG-4MAL hydrogel containing RGD, showing cell-gel interactions through cell spreading and the formation of 3D cell structures. (d-e) Brightfield and fluorescence images of the live/dead assay of cells seeded on top of the bioprinted hydrogels at day 6, showing viable 3D cellular network (green). In a – b, *n* = 3 biological repeats. Scale bars, c = 100 µm and d-e = 200 µm. Results are mean ± SD.

Finally, we assessed the biocompatibility of the bioprinted hydrogel scaffold for its toxicity and biocompatibility with PDAC cells. Prior to printing, RGD attachment to the PEG-4MAL bioink was confirmed using Ellman’s assay by monitoring the disappearance of the signal at 412 nm (Figure S4 (Supporting Information)). Hydrogels comprising 10 wt% PEG-4MAL with 1 mM RGD, crosslinked with bisthiol-PEG were printed onto each well of a 96-well plate. Immediately after printing, 5,000 cells were manually added on top of the hydrogel. After 3 days of incubation, mesenchymal phenotype was visible across the gel surface, consistent with strong integrin-mediated cell-gel interactions (Figure 4c).^[25]^ PDAC on top of the gel showed both cell-gel and cell-cell interactions producing a network-like structure of multiple cells connecting to each other (Figure 4d). Live/dead assay of this sample confirmed a high cell viability, validating that the hydrogels were cell compatible (Figure 4e).

## 4. Bioprinting of 3D Cell Modes with Tunable Biological Properties

To show the effect of the hydrogel microenvironment on cells, we designed two different gels to promote different PDAC phenotypes. PDAC cells carrying both Kras and p53 mutations have been shown previously to be capable of forming both spheroid^[31]^ and mesenchymal morphologies^[32]^ depending on the microenvironment. The first gel (non-cell responsive) was non-MMP degradable and contained no RGD ((f-) RGD/PEG Activator), while the second gel (cell-responsive) was MMP-degradable and decorated with RGD ((+) RGD/MMP Activator). Encapsulated PDAC 3D cell models were prepared by mixing PDAC cells in the activator at 10 million cells/mL. RGD at 2 mM was incorporated into the bioink as it has been previously reported to induce cellular response while still maintaining sufficient crosslinking sites.^[33,34]^ Both the bioink and the cell-activator were primed into the bioprinter and bioprinted into a 96-well plate to form a 3D structure covering the bottom of the wells in a 96-well plate (6.4 mm in diameter, 400 µm thick). This structure was chosen to ensure no interactions between the cells with the tissue culture plastic. After 6 days of incubation, cells in two different hydrogels showed different morphology (**Figure 5**a), while still maintaining > 90% cell viability, quantified via live/dead cell viability analysis (Figure 5b and d).

**Figure 5.**
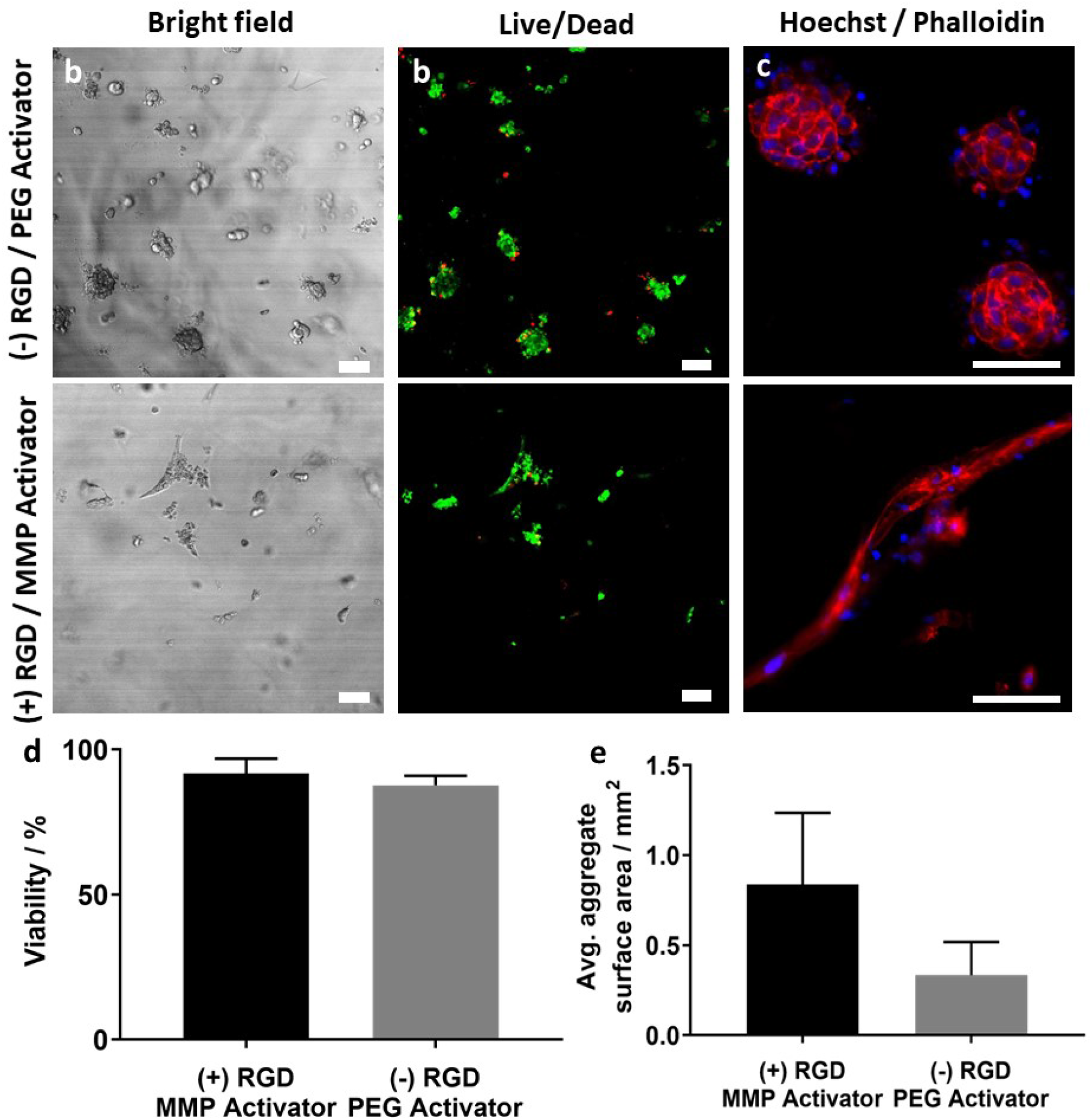
Effect of matrix environment on the morphology of 3D bioprinted pancreatic ductal adenocarcinoma cells assessed after 3 days of incubation. (a) Bright field image of pancreatic cancer cells bioprinted inside a non-MMP responsive hydrogel with no RGD ((-) RGD / PEG Activator) and inside an MMP responsive hydrogel with RGD ((+) RGD / MMP Activator). (b) Live/Dead fluorescence images of the 3D bioprinted cells inside the gel, with their viability confirmed to be > 90% in both cases (d). (e) F-actin/nuclei staining of the 3D bioprinted assay showing the different aggregate morphologies of the cells in two different hydrogel conditions. (e) The difference in morphology was subsequently quantified by measuring the average area of the aggregate, with cells bioprinted in an RGD containing gel forming extensive cellular network and cells bioprinted in non-RGD containing gel conform to a spheroid structure. In d and e, *n* = 3 biological repeats. Scale bars = 200 µm. Results are mean ± SD.

To probe the different cell morphologies further, phalloidin cytoskeleton (red) and Hoechst 33342 nuclei (blue) fluorescence staining were conducted. PDAC in both hydrogels showed the presence of multicellular aggregates as indicated by the presence of multiple cell nuclei (Figure. 5c). In the non-responsive hydrogel system, a multicellular spheroid morphology was dominant, indicating stronger cell-cell interactions amongst the proliferating cells than cell-matrix interactions (Figure 5c, (-) RGD / PEG Activator). Within the cell-responsive hydrogel, multicellular aggregates were showing mesenchymal phenotype with clear spindle protrusions, indicating the presence of cell-gel interactions (Figure 5c, ((+) RGD/MMP-Activator). The quantified surface area of the aggregates further corroborates the finding, showing much tighter aggregate formation occupying a smaller surface area in the non-responsive hydrogel than the aggregates in cell-responsive hydrogel (Figure 5e). The ability of the highly invasive PDAC cells to form large aggregates in the cell-responsive hydrogel also validated the remodeling capability of the hydrogel through cell secreted MMP.

## 5. Conclusion

A multi-material bioprinting platform was developed for high-throughput production of complex 3D cell models. A 4-arm poly(ethylene glycol) maleimide-based hydrogel system was chosen due to its quick gelation mechanism, modularity and cell compatibility. The PEG-4MAL bioink and its complementary activator were optimized to produce accurate and reproducible droplets when printed using the fly-by printing process. Using the fly-by process, various 3D structures comprising three different gel compositions were successful printed. Pancreatic ductal adenocarcinoma cells were chosen to show the modularity and the biocompatibility of the bioprinting platform. The hydrogel and bioprinting process were confirmed to not affect the cell viability. When encapsulated in a cell-responsive hydrogel, elongated multicellular PDAC aggregates with protrusions was visible. In contrast, PDAC spheroids were visible inside the non-cell responsive hydrogel. In both cases cell viability of > 90% was maintained. The combination of multi-material bioprinting capability and biocompatibility would enable the creation of complex, biologically relevant cell models that would be impossible to make manually otherwise.

## 6. Experimental Section

### Materials

4-arm PEG-maleimide (PEG-4MAL, MW 20 kDa, Biochempeg), bisthiol-PEG (MW 1 kDa, Sigma Aldrich), 5,5’-dithiobis(2-nitrobenzoic acid) (Ellman’s reagent, Sigma Aldrich), sodium phosphate (Sigma Aldrich), ethylenediaminetetraacetic acid (Sigma Aldrich), Triton X-100 (Sigma Aldrich), bovine, serum albumin (BSA, Sigma Aldrich), _L_-cysteine hydrochloride monohydrate (Sigma Aldrich), paraformaldehyde (PFA, Electron Microscopy Science), Dulbecco’s phosphate buffer solution (DPBS, ThermoFisher), Dulbecco’s modified eagle’s medium (DMEM, ThermoFisher Scientific), fetal calf serum (FCS, Bovogen), alamarBlue™ (ThermoFisher Scientific), LIVE/DEAD cell viability kit (ThermoFisher Scientific), phalloidin conjugated to Alexa Fluor® 568 (ThermoFisher Scientific), Hoechst 33342 (ThermoFisher Scientific), CRGDS (> 90% purity, Genscript) and GCIPVSLRSGCG (> 90% purity, Genscript) were used as received.

### Bioink Preparation

A bioink of 10 wt% PEG-4MAL bioink was prepared by dissolving 20 kDa PEG-4MAL (0.1 g) in DPBS (1 mL). To prepare an RGD (2 mM) containing PEG-4MAL bioink, CRGDS (1.13 mg) was added into the PEG-4MAL blank bioink (1 mL). The pH of the RGD containing PEG-4MAL bioink solution was then neutralized to pH 7.4 to initiate the RGD coupling reaction. Both solutions were filtered aseptically through a 0.22 µm syringe filter. Sterile solutions were kept at -20 ^°^C prior to use.

### Activator Preparation

All activators were prepared at equimolar thiol concentration to the available maleimide concentration of the complementing bioink. To prepare the bisthiol-PEG activator for 10 wt% PEG-4MAL bioink, bisthiol-PEG (10.5 mg) was dissolved in DPBS (1 mL). To prepare the MMP-responsive activator, GCIPVSLRSGCG (13.8 mg) was dissolved in DPBS (1 mL). The pH of activator was neutralized to pH 7.4 as required. Both solutions were filtered aseptically through a 0.22 µm filter. Sterile solutions were kept at -20 °C prior to use.

### Thiol-maleimide reaction study

The coupling reaction of CRGDS to the maleimide group was monitored using Ellman’s assay. First, Ellman’s assay reaction buffer (pH 7.5 - 8) was prepared by mixing sodium phosphate (0.1 M) and ethylenediaminetetraacetic acid (1 mM). Ellman’s reagent solution was prepared by mixing of 5,5’-dithiobis-(2-nitrobenzoic acid) (4 mg) in the reaction buffer (1 mL). The Ellman’s reagent solution was prepared fresh every time prior to analysis. A 250 µL aliquot of PEG-4MAL alone and PEG-4MAL mixed with CRGDS at 0- and 5-minutes reaction time solution was then mixed with 50 µL of Ellman’s reagent and 2.5 mL of reaction buffer. The solution was incubated at room temperature for 15 minutes, before a UV-Vis absorbance measurement was conducted. To calculate the concentration of free thiol left in the solution, the absorbance value at 412 nm was compared against a calibration curve of _L_-cysteine hydrochloride monohydrate (0 – 1.5 mM).

### Rheology

Rheological property of the hydrogel was measured using an Anton-Paar modular compact rheometer (MCR) 302 equipped with different measuring systems. Viscosity measurements were conducted using the CP-501 measuring system, with a shear rate sweep from 10 – 1000 1/s. The hydrogel storage modulus was measured using the same machine equipped with a parallel-plate geometry (25 mm in diameter, 1 mm thick sample) system and a solvent trap to minimize liquid evaporation. Modulus measurements were conducted on a preheated plate and maintained at a constant temperature of 37 °C. Amplitude strain sweep experiments (*γ*: 1–100%) were carried out to determine the linear viscoelastic region (LVR) at a fixed frequency of 10 Hz. Once the LVR of each sample was established, dynamic frequency sweeps were performed (*ω*: 0.1–10 Hz) at fixed strain in the linear viscoelastic region of the hydrogel. Time sweep measurements were performed at 10 Hz frequency and 0.1% strain amplitude (within the LVR).

### Gelation Time Study

Gel formation was assessed via tube inversion test conducted at room temperature. PEG-4MAL bioink and bisthiol-PEG activator at the desired concentration were initially prepared in a separate vial. Bioink (0.5 mL) was firstly transferred into a 2 mL screwcap, glass vial (OD 12 mm, ThermoFisher Scientific). Subsequently, 0.5 mL of the activator was pipetted in one rapid flow to initiate gelation. The tube was then inverted upside down at the specified time. Gelation was considered complete when no flow was observed when the tube was inverted.

### Cell Culture

Kras^G12D^ and p53^R172H^ mutant pancreatic ductal adenocarcinoma (PDAC) cells were maintained in a Dulbecco’s Modified Eagle’s Medium (DMEM) containing 10% (v/v) fetal calf serum (FCS) at 37 °C and 5% CO_2_. Cell lines are routinely screened and were free of mycoplasma contamination.

### AlamarBlue Cytotoxicity Study

*Pancreatic* ductal adenocarcinoma cells (PDAC) were suspended in the desired bioink or activator at 10 million cells/mL (100 µL final volume). The cell suspension was kept at ambient condition for 30 minutes. Subsequently, 900 µL of 10% (v/v) FCS/DMEM was added to dilute the cell concentration. An aliquot of 5,000 cells from the diluted solution (5 µL) was then transferred into a 96-well plate, topped with 200 µL of culture media and incubated for 3 days. After incubation, the culture media was replaced with alamarBlue™ (200 µL, 10% v/v in 10% (v/v) FCS/DMEM). After 4 h of incubation, the absorbance was measured at 570 nm against 600 nm using a microplate reader. A cell suspension in DPBS, at the same cell concentration was used as the positive control.

### 3D Cell Model Bioprinting

Bioink and activator were allowed to warm to room temperature prior to printing. A cell-laden ink was prepared by mixing 2 million pancreatic ductal adenocarcinoma cells in 200 µL of the activator. Bioprinting was conducted using the RASTRUM modular 3D bioprinter (Inventia Life Science). The structure was designed using the RASTRUM software (Inventia Life Science). The bioprinter was initiated to start the integrated HEPA filtration system and sterilized by ethanol wiping. A sterile cartridge was filled with ethanol (10 mL) and sterile DI water (20 mL). An automated sterilization sequence was followed for the internal fluidics and microvalves. 200 µL bioink and cell-laden activator were then pipetted into their respective reservoirs in the cartridge. Both fluids were then automatically primed into the nozzles.

Each bioprinted sample measured of about 6.4 mm in length and 400 µm in thickness. Both bioink and activator solutions were printed at 30 kPa printing pressure. The generation of the 3D object was achieved by the deposition of a bioink droplet, which then immediately followed by the deposition of an activator droplet on top of it, at the determined location by the software. All printing was conducted in a flat plastic-bottom, 96-well plate (Corning). Upon completion, 150 µL of DMEM, supplemented with 10% (v/v) FCS was added and the plate was transferred into an incubator at 37°C and 5 % CO_2_.

### Multi-materials 3D Bioprinting

Food dyes (10 □ L) were added onto PEG-4MAL bioink. to form solution of different colors (red/green/blue). Bisthiol-PEG activator and colored PEG-4MAL bioink were brought to room temperature prior to printing. Different structures were formulated using the RASTRUM Software (Inventia Life Science) and printed onto each well of a 96-well plate. The resulting structures were captured using Vimicro USB camera.

### Live/Dead Viability Study

Cell viability in 3D cell assays was measured using the LIVE/DEAD cell viability kit containing calcein AM (4 mM stock solution) and ethidium homodimer-I (2 mM stock solution). To prepare 1 mL of the LIVE/DEAD working solution containing 2 µM of calcein AM and 4 µM of ethidium homodimer-1, 0.5 µL of calcein AM and 2 µL of ethidium homodimer-1 stock solution were mixed in 1 mL of 10% (v/v) FCS/DMEM. To stain the bioprinted 3D cell models, the incubating media was replaced with 200 µL of the LIVE/DEAD working solution after 3 d and 6 d of incubation. The samples were further incubated for 2 h at 37 °C/5% CO_2_. The sample was imaged using Leica TCS SP5 confocal microscope at 494 nm and 528 nm to excite calcein AM (green) and ethidium homodimer-1 (red) fluorescence.

### Phalloidin/Hoechst staining

3D bioprinted cell models after 72 h of incubation were fixed, permeabilized and blocked prior to staining. The samples were fixed using 4% paraformaldehyde in DPBS for 30 minutes at 4° C, followed by washing three times in DPBS. The samples were then permeabilized by incubating fixed samples in 0.1% (v/v) Triton X-100/DPBS solution for 20 minutes at 4° C, followed by washing twice with DPBS. Samples were finally blocked by incubation in 2% (v/v) BSA/DPBS for 30 minutes at 37 °C.

Fixed, permeabilized and blocked samples were then incubated in a solution of 2 µL of phalloidin conjugated to Alexa Fluor® 568 stock solution (6.6 μM) diluted in 200 μL 2% (v/v) BSA/DPBS for 1 h. It was then followed by staining with Hoechst 33342 nuclei staining agent, by incubating it in 0.4 μL Hoechst 33342 stock solution stock solution (20 mM) diluted in 200 μL 2% (v/v) BSA/DPBS for 30 minutes. The stained sample was washed once with PBS and imaged using a Zeiss LSM 780 confocal microscope at 578 nm and 455 nm for the phalloidin and Hoechst 33342 fluorescence, respectively.

### Image Acquisition and Analysis

High magnification images of the 3D bioprinted hydrogel were acquired using a custom-built plate imaging system (Inventia Life Science). The system was equipped with a Bi-telecentric lens (TC23016, Opto Engineering) connected to a Mako G-192 (Adept Turnkey) industrial camera. Acquired images were saved as PNG.

LIVE/DEAD brightfield and fluorescence images were acquired using Leica TCS SP5 (Leica camera AG) on an inverted configuration. All images were obtained using a 20x objective lens, NA 0.5, without immersion. A Hybrid Detector (HyD) was used. Acquired images were saved in the native LIF format and were processed and analyzed using ImageJ 1.50 software.

Immunofluorescence images were acquired using Zeiss LSM 780 (Carl Zeiss AG) operating in the inverted configuration. All images were obtained using a 40x magnification objective lens, NA 0.65, without immersion. Both Hamamatsu blue sensitive PMT and gallium arsenide phosphide (GaAsP) photodetector were used to detect below and above 450 nm wavelength, respectively. Images saved in the native CZI format were processed and analyzed using ImageJ 1.50 software.

### Statistical analysis

Statistical analyses were performed using the GraphPad Prism v8.1.0 software (GraphPad Software). Results were expressed as means of the number of independent repeats performed ± standard deviation (SD).

## Acknowledgements

The authors would like to thank the Biomedical Imaging facility (BMIF) within the Mark Wainwright Analytical Centre at UNSW for the use of the facility and their technical assistance. The authors acknowledge the following funding sources: The Australian Centre for NanoMedicine (ACN) Seed Funding (R.H.U., V.T.G.T); The Innovation Connection Grant (R.H.U., J.J.G.); the Australian Research Council (ARC) Linkage Grant (LP170100623 to J.J.G., M.K., J.R.); ARC Laureate Fellowship (FL150100060 to J.J.G); National Health and Medical Research Council (NHMRC) Program Grant (APP1091261 to M.K, J.J.G); NHMRC Principal Research Fellowship (APP1119152 to M.K), NHMRC Investigator (APP1196648 to J.J G.) and grant support from ARC Centre of Excellence in Convergent Bio-Nano Science and Technology (CE140100036 to J.J.G and M.K). We would like to thank Prof. Paul Timpson for providing us with the PDAC cells.

