## Supplementary Information for "A covalently crosslinked bioink for multi-materials drop-on-demand 3D bioprinting of three-dimensional cell cultures"

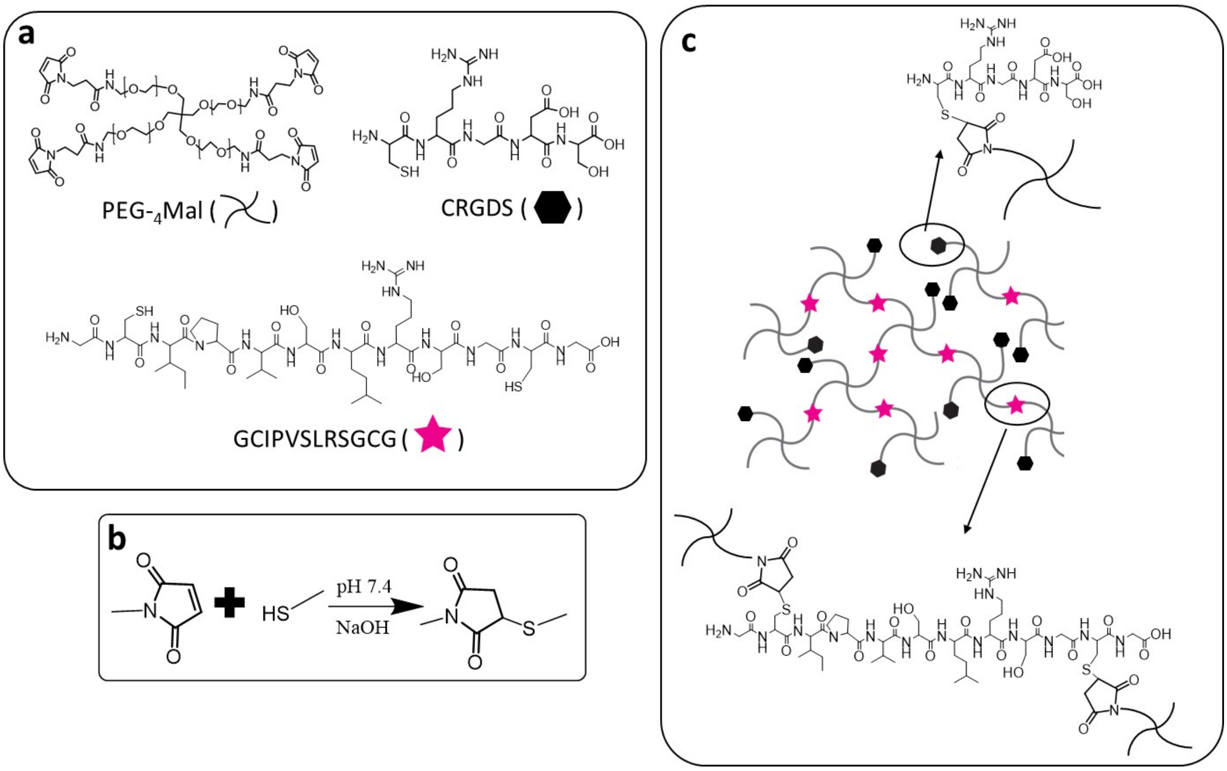


**Figure S1.** PEG-4MAL hydrogel formation chemistry. (a) (+) RGD / MMP Activator PEG-4MAL hydrogel components comprising PEG-4MAL polymer, CRGDS adhesion peptide and GCIPVSLRSGCG MMP-responsive crosslinker. (b) A reaction schematic of thiol-Michael addition. (c) A schematic representation of the (+) RGD / MMP Activator PEG-4MAL hydrogel network with both RGD attachment and MMP-activator crosslinking facilitated by thiol-Michael addition reaction.

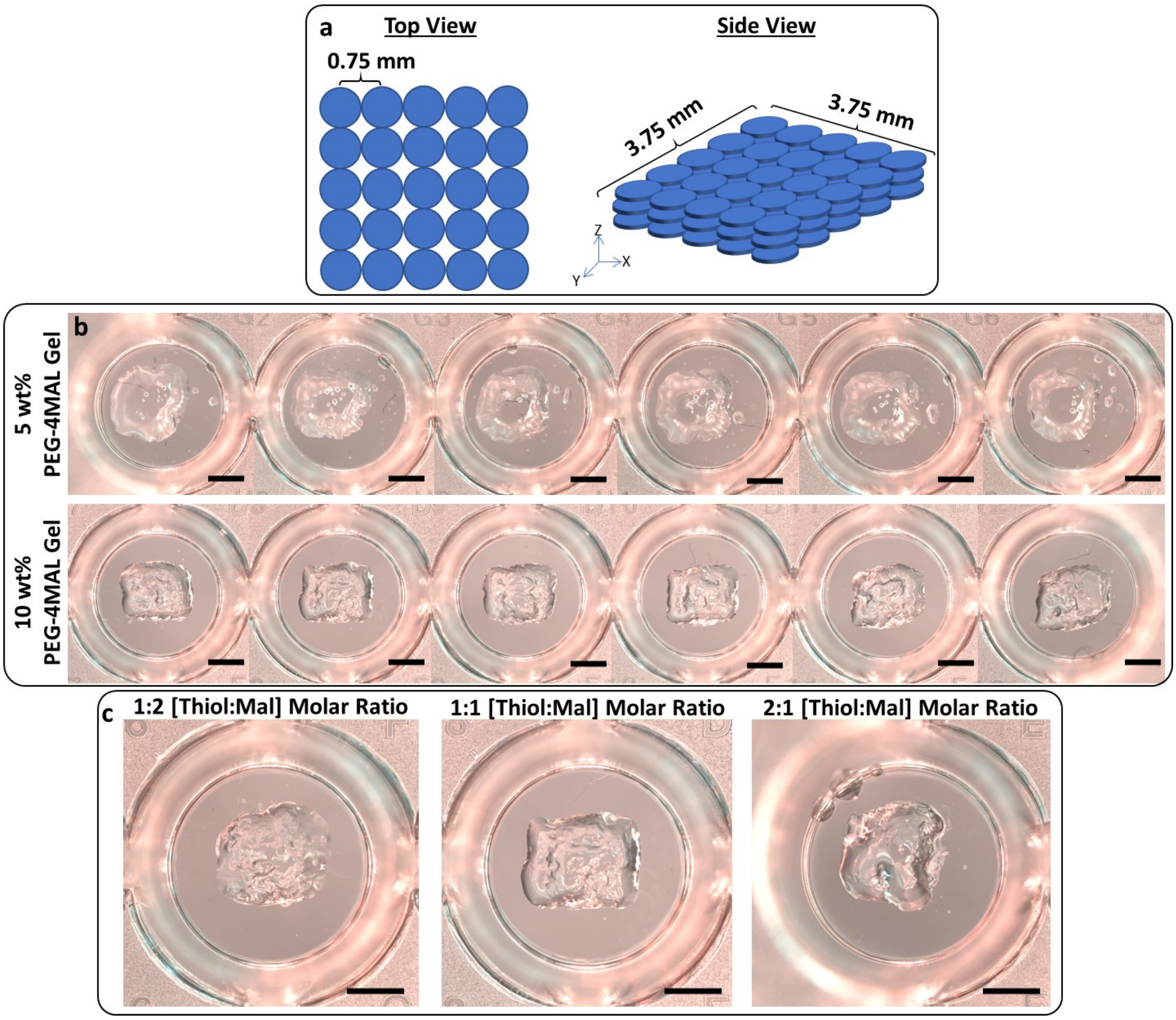


**Figure S2.** Assessment of various gelation properties on the 3D bioprinted structure fidelity. (a) Schematic diagram of the designed 3D structure. High magnification images of a 5x3-droplet, 3-layer structure printed on a 96-well plate using (b) PEG-4MAL bioink at 5 wt% and 10 wt% concentration and bisthiol-PEG activator at 1:1 thiol:maleimide (Thiol:Mal) molar ratio and (c) PEG-4MAL bioink at 10 wt% concentration and bisthiol-PEG activator at 1:2, 1:1 and 2:1 thiol:maleimide molar ratio. Scale bars = 2 mm

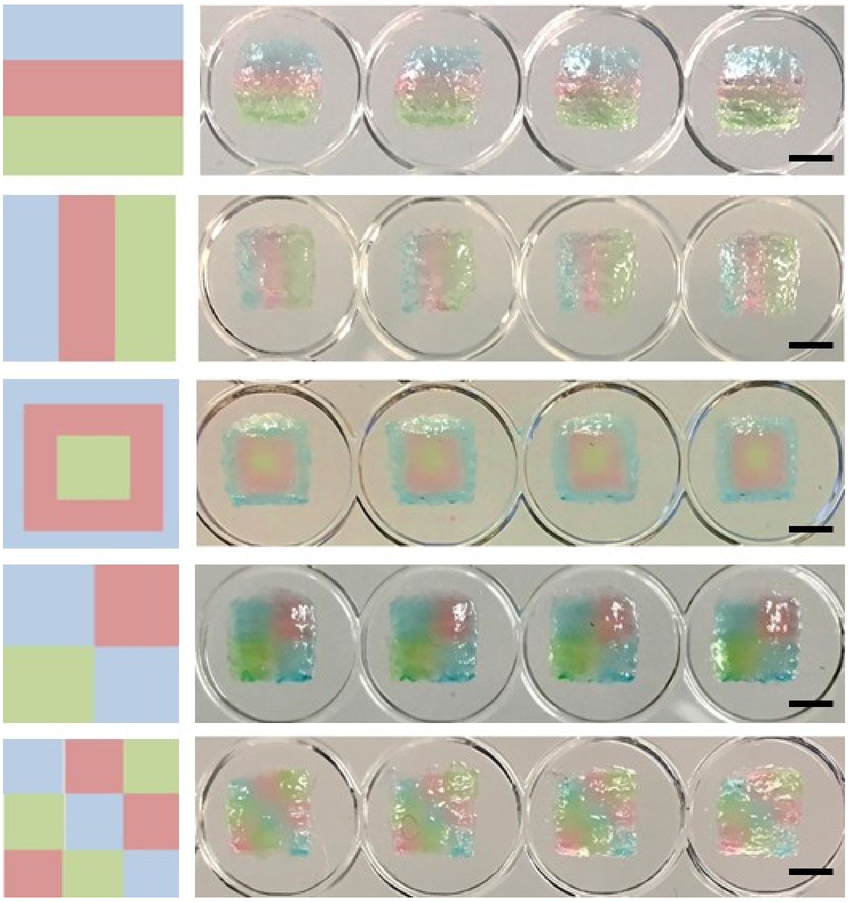

**Figure S3**. Multi-material printing of colored 4-arms PEG hydrogel of multiple samples, showing the consistency of the system in printing complex 3D structures. Scale bars = 2 mm

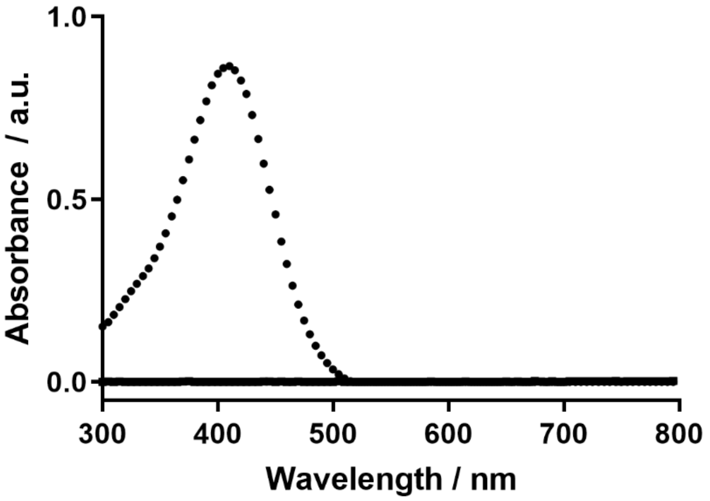


**Figure S4.** UV-Vis spectrum of the bioink solution of PEG-4MAL with CRGDS at 0 minute (circle) and 5 minutes (square). The signal at 412 nm was used to quantitatively calculate the amount of thiol (from CRGDS) left in the solution.

**Table S1**: Effect of PEG-4MAL bioink concentration on gelation time, tested using tube-inversion test.

| PEG-4MAL Bioink [wt%] | Gelation time [second] |
|---|---|
| 10 | < 1 |
| 5 | 3 |
| 3 | 10 |
| 2 | 62 |
| 1 | >300 |

**Table S2:** Comparison of various thiol:maleimide molar concentration ratios and its effect on gelation properties. 10 wt% PEG-4MAL 20 kDa bioink was used in all cases. The concentration of bisthiol-PEG was altered to reflect the three different ratios.

| Molar thiol-maleimide ratio | Observation |
|---|---|
| 2:1 | Soft and flowing gel under gravity formed after 1 minute |
| 1:1 | Instantaneous gelation (< 1 second) upon mixing |
| 1:2 | Instantaneous gelation (< 1 second) upon mixing |

**Table S3**: The composition of the bioink and activator for each PEG-4MAL hydrogel concentration tested.

| Hydrogel concentration [wt%] | PEG-4MAL concentration in bioink [wt%] | Bis-thiol peptide concentration in activator [wt%] |
|---|---|---|
| 5 | 10 | 1.26 |
| 10 | 20 | 2.52 |
| 15 | 30 | 3.78 |
| 20 | 40 | 5.04 |
